# Identification and qualification of 500 nuclear, single-copy, orthologous genes for the Eupulmonata (Gastropoda) using transcriptome sequencing and exon-capture

**DOI:** 10.1101/035543

**Authors:** Luisa C. Teasdale, Frank Köehler, Kevin D. Murray, Tim O’Hara, Adnan Moussalli

**Affiliations:** Sciences Department, Museum Victoria, 11 Nicholson St, Carlton, Vic, Australia 3053; School of BioSciences, The University of Melbourne, Parkville, Vic, Australia 3010; Australian Museum, 6 College Street, Sydney, NSW, Australia 2010; Division of Plant Sciences, Research School of Biology, Australian National University, Australia 0200

**Keywords:** orthology, single-copy, phylogenomics, Pulmonata, transcriptome, targeted enrichment

## Abstract

The qualification of orthology is a significant challenge when developing large, multiloci phylogenetic datasets from assembled transcripts. Transcriptome assemblies have various attributes, such as fragmentation, frameshifts, and mis-indexing, which pose problems to automated methods of orthology assessment. Here, we identify a set of orthologous single-copy genes from transcriptome assemblies for the land snails and slugs (Eupulmonata) using a thorough approach to orthology determination involving manual alignment curation, gene tree assessment and sequencing from genomic DNA. We qualified the orthology of 500 nuclear, protein coding genes from the transcriptome assemblies of 21 eupulmonate species to produce the most complete gene data matrix for a major molluscan lineage to date, both in terms of taxon and character completeness. Exon-capture targeting 490 of the 500 genes (those with at least one exon > 120 bp) from 22 species of Australian Camaenidae successfully captured sequences of 2,825 exons (representing all targeted genes), with only a 3.7% reduction in the data matrix due to the presence of putative paralogs or pseudogenes. The automated pipeline Agalma retrieved the majority of the manually qualified 500 single-copy gene set and identified a further 375 putative single-copy genes, although it failed to account for fragmented transcripts resulting in lower data matrix completeness. This could potentially explain the minor inconsistencies we observed in the supported topologies for the 21 eupulmonate species between the manually curated and Agalma-equivalent dataset (sharing 458 genes). Overall, our study confirms the utility of the 500 gene set to resolve phylogenetic relationships at a broad range of evolutionary depths, and highlights the importance of addressing fragmentation at the homolog alignment stage for probe design.

## INTRODUCTION

Robust and well resolved phylogenies document the evolutionary history of organisms and are essential for understanding spatio-temporal patterns of phylogenetic diversification and phenotypic evolution. Despite the central role of phylogenies in evolutionary biology, most phylogenetic studies in non-model systems have relied on a limited number of readily sequenced genes due to cost restrictions and availability of phylogenetic markers. However, both theoretical and empirical studies have shown that a greater number of independently evolving loci are needed to resolve difficult phylogenetic questions (Gontcharov *et al*. 2004; Wortley *et al*. 2005; Leaché & Rannala 2011). This need has been addressed by rapid advances in phylogenomics, which capitalise on high-throughput sequencing to acquire large multi-loci datasets. In particular, both transcriptome sequencing and targeted-enrichment strategies are increasingly being employed to reconstruct phylogenetic relationships across a wide range of taxonomic levels (e.g. Bi *et al*. 2012; Lemmon *et al*. 2012; Faircloth *et al*. 2012; Zapata *et al*. 2014; O’Hara *et al*. 2014; Misof *et al*. 2014). A common aim of these studies, especially targeted enrichment based studies, has been to identify universal sets of orthologous loci that can readily be captured and sequenced across a broad taxonomic spectrum (e.g. Lemmon *et al*. 2012; Faircloth *et al*. 2012; Hugall *et al*. 2015). Obtaining such universal sets of orthologous genes allows for consistency and comparison across studies, and ultimately contributes towards a more comprehensive Tree of Life (ToL) meta-analysis.

One of the greatest challenges associated with developing large, multi-loci phylogenomic datasets is the qualification of orthology. In the context of phylogenetic analysis, genes need to be orthologous and single-copy across all taxa under study (Fitch 2000; Philippe *et al*. 2011; Struck 2013). To this end, a number of automated pipelines have been developed to identify single-copy orthologous genes from assembled transcriptomes. These methods generally involve two main steps. The first step is to identify and cluster homologous sequences, either by direct reference to annotated genomes (e.g., O’Hara *et al*. 2014) or by reference to ortholog databases, which themselves are derived from genome comparisons (e.g., Tatusov *et al*. 2003; Ranwez *et al*. 2007; Waterhouse *et al*. 2013; Altenhoff *et al*. 2015). Alternatively, non-reference methods have been employed such as all-by-all and reciprocal BLAST comparisons (Li *et al*. 2003; Dunn *et al*. 2013) followed by clustering (Enright *et al*. 2002). In the second step, orthology is qualified using either similarity based approaches, including best-hit reciprocal blasts (Ebersberger *et al*. 2009; Waterhouse *et al*. 2013; Ward & Moreno-Hagelsieb 2014), and/or tree based methods, where gene trees are used to identify sequences with purely orthologous relationships (e.g., Agalma, Dunn *et al*. 2013; PhyloTreePruner, Kocot *et al*. 2013; TreSpEx, Struck 2014).

Despite rapid advances in automated approaches to homolog clustering and qualifying orthology, there are many characteristics of transcriptome assemblies that challenge such automated methods. These include frameshifts, mis-indexing, transcript fragmentation and the presence of multiple isoforms. Not accounting for these issues can lead to erroneous inclusion of paralogous sequences and/or the inadvertent removal of appropriate orthologous sequences (Martin & Burg 2002; Pirie *et al*. 2007; Philippe *et al*. 2011). To address these issues O’Hara *et al*. (2014) placed greater emphasis on careful manual curation and editing of homolog alignments prior to orthology qualification. A key aspect of this approach was the concatenation of transcript fragments into a single consensus sequence prior to tree-based ortholog qualification, leading to a more complete final data matrix. This, in turn, allowed a more robust probe design for subsequent exon-capture (Hugall *et al*. 2015). With the same objective of deriving a gene set appropriate for exon-capture in future studies, here we implement this approach to identify and qualify 500 single-copy orthologous genes for the Eupulmonata, a major lineage of air breathing snails and slugs within the class Gastropoda.

Eupulmonata comprises over 20,000 species, with an evolutionary depth spanning over 150 million years (Jörger *et al*. 2010; Lydeard *et al*. 2010). The evolutionary relationships of the Eupulmonata, however, remain incompletely understood despite many morphological and molecular phylogenetic studies over the last two decades (e.g., Ponder & Lindberg 1997; Wade *et al*. 2001, 2006; Grande *et al*. 2004; Dinapoli & Klussmann-Kolb 2010; Holznagel *et a1*. 2010; Dayrat *et al*. 2011). The lack of congruence between studies is largely due to a combination of using insufficient genetic markers (Schrödl 2014), with many studies relying on 28S rRNA or mitochondrial sequences, and widespread morphological convergence (Dayrat & Tillier 2002). Therefore to resolve the ‘tree of life’ of the eupulmonates, it is essential to identify more independently evolving markers, with a greater range of substitution rates, to better estimate relationships across all evolutionary depths. To achieve this, we sequenced and assembled transcriptomes for representatives of 15 families across Eupulmonata. We used the owl limpet genome, *Lottia gigantea*, as a reference to identify and cluster homologous sequences and visually assessed and manually edited candidate homolog alignments accounting for transcript fragmentation, mis-indexing and frameshifts. We then further qualified orthology by assessing individual gene trees and by sequencing the orthologous gene set from genomic DNA using exon-capture as unexpressed paralogs or pseudogenes will not be detected in transcriptome datasets. Lastly, as a comparison and qualification of our approach we also analysed our transcriptome dataset using the fully automated orthology determination pipeline Agalma (Dunn *et al*. 2013).

## METHODS

### Transcriptome sequencing and assembly

We sequenced transcriptomes for 21 species of terrestrial snails and slugs representative of 15 families across Eupulmonata (Table 1). Total RNA was extracted from foot or whole body tissue stored in RNAlater (Ambion Inc, USA) using the Qiagen RNeasy extraction kit (Qiagen, Hilden, Germany). Library preparations were conducted using the TruSeq RNA sample preparation kit v2 (Illumina Inc., San Diego, CA), and sequenced on the Illumina HiSeq 2000 platform (100 bp paired end reads). We used the program Trimmomatic v0.22 (Lohse *et al*. 2012) to remove and trim low quality reads and adaptor sequences, and the program Trinity v2012-06-08 (Grabherr *et al*. 2011; Haas *et al*. 2013) with default settings to assemble the transcriptomes.

**Table 1.** Taxon sampling: Transcriptome sequencing

| Superfamilies or higher unranked classification | Family | Species | Voucher specimen | Collection locality* |
| --- | --- | --- | --- | --- |
| Helicoidea | Camaenidae | *Austrochloritis kosciuszkoensis* Shea & Griffiths, 2010 | NMV F193285 | Sylvia Creek, VIC |
| Helicoidea | Camaenidae | *Chloritobadistes victoriae* (Cox, 1868) | NMV F193288 | Crawford River, VIC |
| Helicoidea | Camaenidae | *Ramogenia challengerii* (Gude, 1906) | NMV F193287 | Noosa, QLD |
| Helicoidea | Camaenidae | *Sphaerospira fraseri* (Griffith & Pidgeon, 1833) | NMV F193284 | Noosa, QLD |
| Helicoidea | Camaenidae | *Thersites novaehollandiae* (Gray, 1834) | NMV F193248 | Comboyne, NSW |
| Helicoidea | Helicidae | *Helix aspersa* Müller, 1774 | NMV F193280 | Melbourne, VIC |
| Limacoidea | Dyakiidae | *Asperitas stuartiae* (Pfeiffer, 1845) | NMV F193286 | North of Dili, Timor-Leste |
| Limacoidea | Helicarionidae | *Fastosarion cf virens* (Pfeiffer, 1849) | NMV F193282 | Noosa, QLD |
| Limacoidea | Limacidae | *Limax flavus* Linnaeus, 1758 | NMV F193283 | Melbourne, VIC |
| Limacoidea | Microcystidae | *Lamprocystis* sp. | AM C.476947 | Ramelau Mountains, Timor-Leste |
| Limacoidea | Milacidae | *Milax gagates* (Draparnaud, 1801) | NMV F226625 | Melbourne, VIC |
| Limacoidea | Oxychilidae | *Oxychilus alliarius* (Miller, 1822) | NMV F226626 | Melbourne, VIC |
| Orthurethra | Cerastidae | *Amimopina macleayi* (Brazier, 1876) | NMV F193290 | Darwin, NT |
| Orthurethra | Cochlicopidae | *Cochlicopa lubrica* (Müller, 1774) | MV614 | Blue Mountains, NSW |
| Orthurethra | Enidae | *Apoecus apertus* (Martens, 1863) | AM C.488753 | Ramelau Mountains, Timor-Leste |
| Rhytidoidea | Rhytididae | *Austrorhytida capillacea* (Férussac, 1832) | NMV F193291 | Blue Mountains, NSW |
| Rhytidoidea | Rhytididae | *Terrycarlessia turbinata* Stanisc, 2010 | NMV F193292 | Comboyne, NSW |
| Rhytidoidea | Rhytididae | *Victaphanta atramentaria* (Shuttleworth, 1852) | NMV F226627 | Toolangi, VIC |
| Ellobioidea | Ellobiidae | *Cassidula angulifera* (Petit, 1841) | NMV F193289 | Manatuto, Timor-Leste |
| Otinoidea | Smeagolidae | *Smeagol phillipensis* Tillier & Ponder, 1992 | MVR13_138 | Phillip Is., VIC |
| Veronicelloidea | Veronicellidae | *Semperula maculata* (Templeton, 1858) | AM C.476934 | Manatuto, Timor-Leste |

\*All localities within Australia unless otherwise indicated

### Homolog clustering

Our approach to homolog clustering and orthology qualification is largely consistent with that detailed in O’Hara *et al*. (2014). A schematic representation of our pipeline is provided in Figure 1. First, to generate clusters of putatively homologous sequences we compared each assembly to the *Lottia gigantea* predicted gene dataset (hereon referred to as the *L. gigantea* genes). The *L. gigantea* reference represents 23,851 filtered gene models annotated in the most current draft genome (Grigoriev *et al*. 2012; Simakov *et al*. 2013). Each transcriptome assembly was compared against the *L.gigantea* genes using blastx with an e-value cut off of 1e-10. This is a relatively relaxed threshold given the small size of the *L. gigantea* reference set. A relaxed e-value cutoff was used to ensure all closely related homologs were assessed without allowing through too many spurious matches with non-homologous sequences. We retained only the top hit for each assembled contig (i.e. the match with the lowest e-value).

**Fig. 1.**
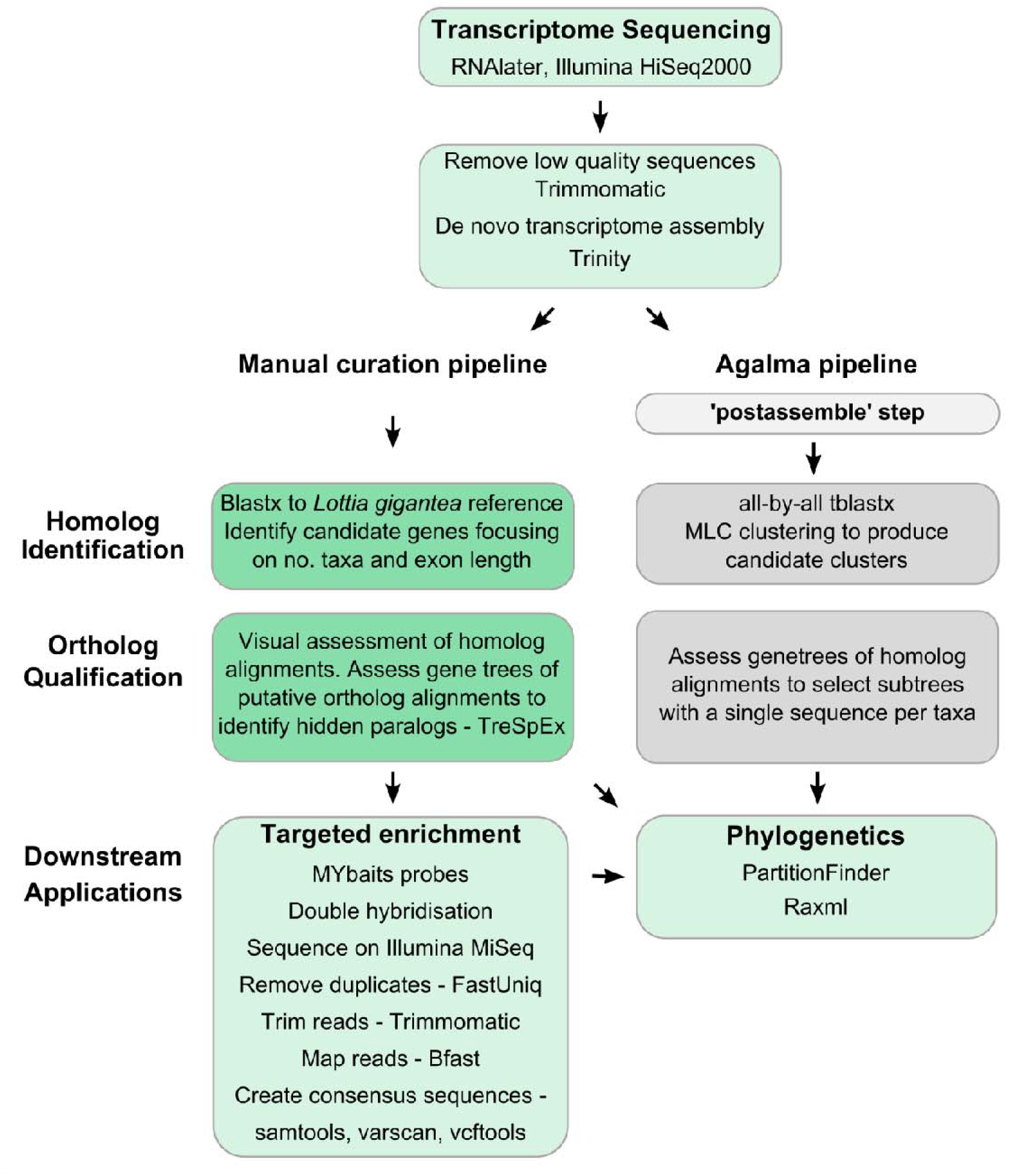
Analysis Pipelines. Outline of the two pipelines used to detect single-copy, orthologous genes from 21 eupulmonate transcriptomes.

In addition to identifying homologous contigs from each transcriptome assembly, we also identified putative paralogs within the *L. gigantea* genome itself, in order to aid the identification of paralogous sequences within the eupulmonates. We ran an all-by-all BLAST of the *L. gigantea* genes against themselves (blastp, cut off e-value of 1e-10), retaining all hits to identify *L. gigantea* genes which had hits to *L. gigantea* genes other than themselves. To qualify the all-by-all BLAST results, we also obtained orthology status for all *L. gigantea* genes classified in the Orthologous MAtrix (OMA) ortholog database (Altenhoff *et al*. 2015). A *L. gigantea* gene was considered to be single-copy if it was the only *L. gigantea* sequence in its respective OMA group. While this information provided guidance, we were not reliant on the *L. gigantea* orthology status when prioritising homolog clusters to assess (see below for criteria used). We considered *L. gigantea* to be sufficiently divergent from the eupulmonates (> 400 million years, Zapata *et al*. 2014) that single-copy status could differ.

The BLAST results for both the transcriptomes compared to *L. gigantea* and the *L. gigantea* all-by-all BLAST were used to produce clusters of homologous sequences linked by having a match to a specific *L. gigantea* gene. Hence, a homolog cluster represents 1) all contigs from all species transcriptomes that had a BLAST match to a given reference *L. gigantea* gene (there were often multiple contigs per taxon with hits to a given *L. gigantea* gene), and 2) all contigs having a hit to any of the closely related *L. gigantea* genes identified by the all-by-all BLAST.

### Orthology assessment

After constructing the homologous clusters, we first visually assessed the alignments for evidence of paralogy. Sequences for each cluster were placed into the correct reading frame using coordinates output from the Blastx comparison for each transcriptome against *L. gigantea*, and were then translated and aligned in amino acids using ClustalW (Thompson *et al*. 1994) within the program BioEdit (Hall 1999). We only considered the coding region (i.e. untranslated regions (UTRs) were removed) which was identified manually by reference to the *L. gigantea* protein sequence for the relevant gene, which was included in the alignments. Many of the homolog clusters contained multiple fragmented transcripts for a given species that were shorter than the coding region but which often overlapped. These fragmented transcripts were synthesised into consensus sequences by manual manipulation within BioEdit, if the overlapping regions did not differ by more than three nucleotides. Nonoverlapping fragments were also concatenated if there were no competing contigs covering the same region of the alignment and both sequences displayed a high degree of similarity to non-fragmented sequences in closely related taxa.

By visually assessing each homolog alignment in both amino acid and nucleotides (in Bioedit it is straight forward to toggle between the two), we were able to identify and manually correct frameshifts. These were clearly evident as a large proportion of a contig would not align with the rest of the sequences and the site of the frameshift was usually associated with runs of adenines. We also manually edited the alignments to remove clearly erroneous sequences which could not be aligned, clear out-paralogs (i.e. sequences which are paralogous but the duplication event took place before the common ancestor of eupulmonates) and redundant sequences (identical transcripts within a species). Mis-indexing was identified as cases where, within the one assembly, two contigs were present for the same region but one (typically the shorter contig having low coverage) matched the sequence for another taxon exactly. Taxa containing paralogs were clearly evident in the alignments as they frequently had > 5% dissimilarity at the nucleotide level between overlapping contigs within the one sample. To further qualify that these sequences were paralogs we inspected genealogies constructed using the neighbour joining method in MEGA (see Figure S1). Any homolog cluster containing paralogs for any species was excluded from further consideration. In certain cases paralogous sequences were closely related (3-5% dissimilarity), representing either in-paralogs (see Remm *et al*. 2001) or genes exhibiting elevated allelic diversity (see O’Hara *et al*. 2014). These genes were also excluded from further consideration as such genes are not optimal for exon-capture.

Approximately 1,500 homologous clusters were visually assessed in order to find 500 which were orthologous across all 21 taxa assessed. This dataset size was chosen to represent a balance between phylogenetic power at varying time scales (Leaché & Rannala 2011; Philippe *et al*. 2011; Lemmon & Lemmon 2013) and a suitable size for subsequent exon-capture probe design. To maintain consistency across studies, we first assessed homolog alignments corresponding to the 288 *L. gigantea* genes used in a phylogenomic study of the Mollusca (Kocot *et al*. 2011). Although there are two other published molluscan phylogenomic datasets (Smith *et al*. 2011; Zapata *et al*. 2014), we focussed on the final dataset of Kocot *et al*. (2011) as the *L. gigantea* gene IDs were documented in the supplementary they provided, which in turn allowed us to easily identify and assess these genes given our pipeline was based on the same reference. We then proceeded to assess and qualify additional homolog clusters until we obtained a final set of 500 single-copy orthologous genes. Accordingly, we prioritised homolog clusters with high taxonomic representation (≥ 18 taxa), as completeness of the data matrix is critical for designing probes across multiple lineages (Lemmon *et al*. 2012; Hugall *et al*. 2015). Where possible we also prioritised homolog alignments for which the corresponding *L. gigantea* gene had a coding region (CDS) ≥ 300 bp or had at least one exon ≥ 200 bp.

As a proxy measure of substitution rate variation across the final 500 gene set, we calculated uncorrected distances (p-distance) for species pairs within the families Rhytididae (*Terrycarlessia turbinata* and *Victaphanta atramentaria*) and Camaenidae (*Sphaerospira fraseri* and *Austrochloritis kosciuszkoensis*). We chose to limit this analysis to intrafamilial comparisons to avoid underestimation due to saturation. For comparison, we also calculated the p-distances for two commonly used phylogenetic markers, CO1 and 28S, for the same taxa.

### Qualification of orthology using gene tree assessments

Although only a single copy of each gene per taxon was present in our final ortholog alignments, they may nevertheless be paralogous across taxa (see Struck 2014). To investigate ‘hidden paralogy’ we used the program TreSpEx (Struck 2014) to assess genealogies for conflict with *a priori* taxonomic hypotheses. Gene trees for each of the 500 genes were constructed using the GTRGAMMA model, codon specific partitioning, and 100 fast bootstraps in RAxML (Stamatakis 2006). TreSpEx then identified well supported conflicting phylogenetic signal relative to five distinct and taxonomically well-established eupulmonate clades (Limacoidea, Orthurethra, Helicoidea, the Australian Rhytididae (Table 1: see Hausdorf 1998; Wade *et al*. 2006, 2007; Herbert *et al*. 2015), and the Stylommatophora). All nodes with ≥ 75 bootstrap support were first assessed for conflict with the monophyly of each of the five clades. Strongly supported sister relationships between sequences from different clades can indicate the presence of ‘hidden’ paralogous sequences. TreSpEx flags very short terminal branches (parameter blt set to 0.00001) as indicative of potential cross-contamination and internal branches which are five times greater than the average (parameter lowbl set to 5), which, in addition to strong nodal support, may indicate paralogy.

### Qualification of orthology using exon-capture

To further qualify orthology and identify unexpressed paralogs and pseudogenes, we designed an exon-capture probe set to enrich and sequence exons from our 500 gene dataset. As the divergence across the eupulmonates is too large for a single probe design we designed a probe set for the Australian Camaenidae as a test case. It would be feasible, however, to design a probe set from our alignments for any of the taxa we have assessed in this study. We designed the baits based on two species of Australian Camaenidae, *Sphaerospira fraseri* and *Austrocholritis kosciuszkoensis*, which represent two divergent lineages of the Australian camaenids (Hugall & Stanisic 2011). Specifically, we included sequences from both taxa for each gene in the probe design. The divergence between these taxa ranges up to ~12% (Figure 3) which is about the level of divergence tolerated be the probes (Hugall *et al*. 2015). Including both taxa in the design increases the likelihood that we will capture sequences from more divergent lineages within the Camaenidae for which we don’t yet have transcriptome sequences. Exon boundaries were first delineated using the program Exonerate v2.2.0 (Slater & Birney 2005) in reference to the *L. gigantea* genomic sequences and then manually qualified using the boundaries detailed in the *L. gigantea* genome annotation (JGI, Grigoriev *et al*. 2012). All exons shorter than 120bp (the probe length) were excluded. This resulted in a target consisting of 1,646 exons from 490 of the 500 genes (ten genes contained only exons shorter than 120bp and were excluded from the bait design). Probes for the target sequences were designed and produced by MYcroarray (Ann Arbor, Michigan) using MYbaits custom biotinylated 120bp RNA baits at 2X tiling.

**Figure 2.**
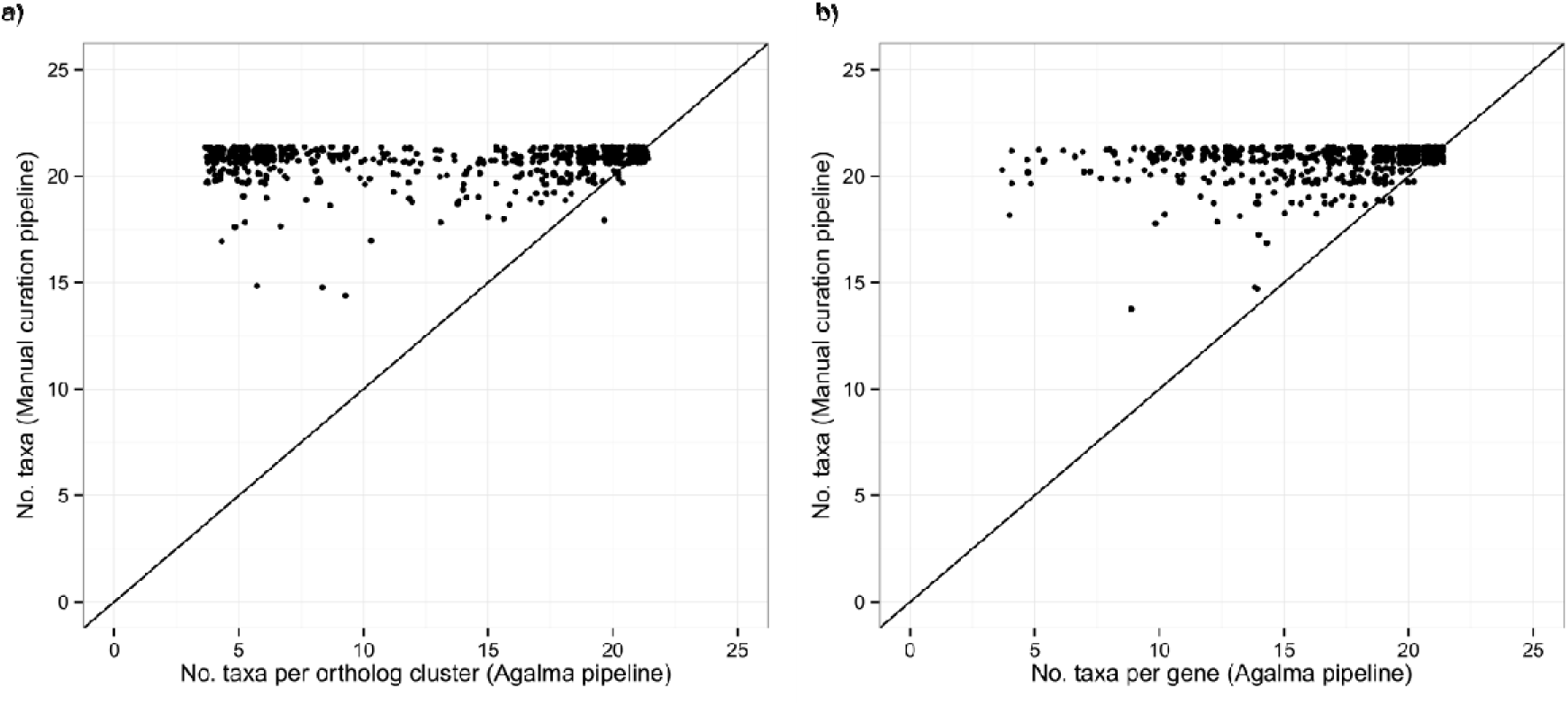
A comparison between two orthology detection pipelines. (a) shows the relationship between the number of taxa per ortholog cluster for the ortholog clusters in common between the manual curation and Agalma pipelines. The manually curated alignments resulted in more taxa complete alignments than the corresponding Agalma alignments.(b) shows the same relationship, however, the number of taxa per gene for the Agalma pipeline were calculated across all ortholog clusters which matched the same *L. gigantea* gene. A comparison of the two plots demonstrates that Agalma tended to produce multiple independent alignments per *L. gigantea* gene, whereas a single alignment was produced through manual curation. Even when the number of taxa recovered across all Agalma alignments associated with a given gene are summed, taxa completeness of the Agalma dataset remained lower than that obtained through manual curation (see also Figure 4e). These graphs are plotted using geom_jitter in ggplot2 to help visualise the large number of data points.

**Figure 3.**
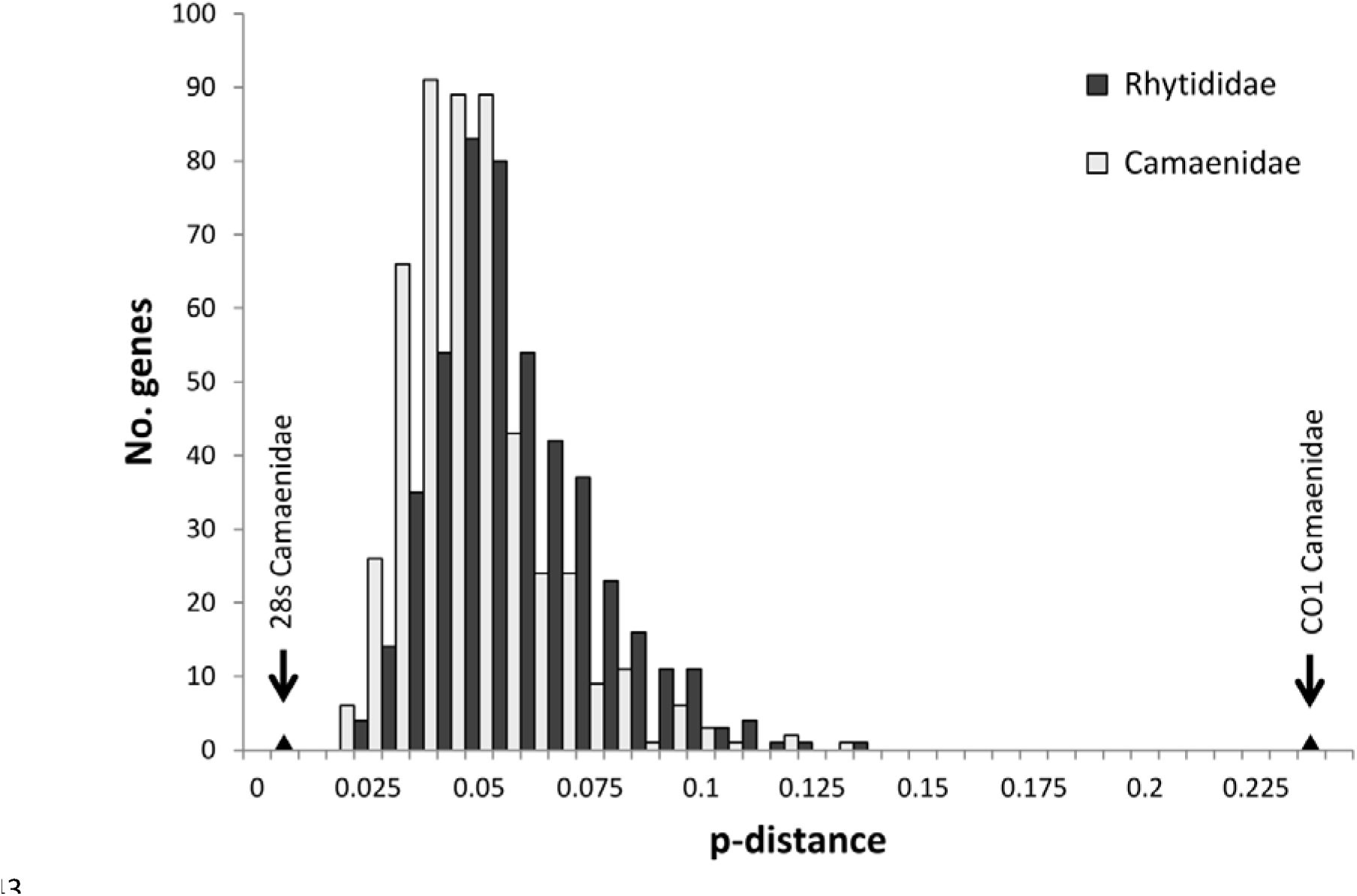
Distribution of the p-distance for 500 single-copy orthologous genes across two families. Uncorrected distances for both groups were calculated using alignments of *Terrycarlessia turbinata* and *Victaphanta atraamentaria* (Rhytididae), and *Austrochloritis kosciuszkoensis* and *Sphaerospira fraseri* (Camaenidae). Triangles on the x-axis notate p-distances of two commonly used phylogenetic markers, CO1 and 28S, for the Camaenidae.

**Figure 4.**
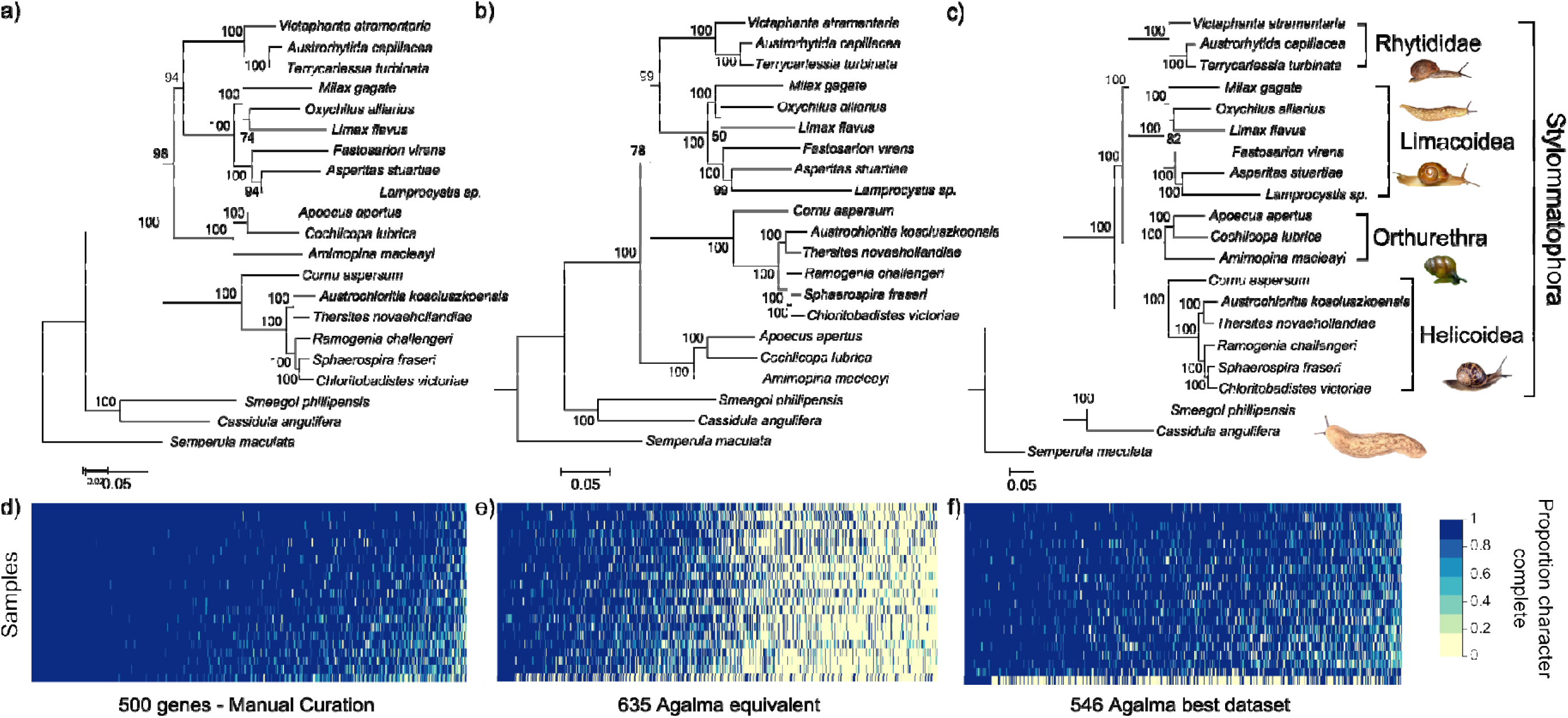
Maximum Likelihood phylogenies for 21 eupulmonates based on three datasets. These datasets were (a) 500 nuclear single-copy orthologous genes identified by manual curation, (b) 635 orthologous clusters identified by the automated pipeline Agalma, which correspond the same 500 genes, and (c) 546 orthologous clusters identified by Agalma, where each orthologous cluster was the only one produced from the respective homolog cluster and had sequences for at least 18 taxa. Phylogenies are each based on analyses of amino acid sequences. Numbers on branches indicate bootstrap nodal support. Heat maps (d, e, f) indicate proportions of sequence obtained for each gene per sample for each dataset (sorted left to right by total proportion of data present per gene, top to bottom by total proportion of data present per sample).

We tested the probe set on 22 camaenid species spanning much of the phylogenetic breadth of the Australasian camaenid radiation, representing up to 30 million years (My) of evolution (Hugall & Stanisic 2011) (Table 2). DNA was extracted using the DNeasy blood and tissue kit (Qiagen) and sheared using the Covaris S2 (targeting a fragment size of 275bp). Libraries where then constructed using the Kapa DNA Library Preparation Kit (Kapa Biosystems, USA), modified to accommodate dual-indexing using the i7 and i5 index sets (see Hugall *et al*. 2015). Up to eight libraries (normalised to 100 ng each) were pooled per capture, and hybridised to the baits (at one-quarter dilution) for 36 hours, following the MYbait protocol v1. A second hybridisation was then carried out on the fragments retained from the first hybridisation to further enrich the capture. Several captures were then multiplexed and sequenced on the Illumina MiSeq platform (v2), obtaining 150bp paired-end reads.

**Table 2.** Taxon sampling: Transcriptome sequencing

| Species | Voucher specimen | Collection locality* |
| --- | --- | --- |
| *Boriogenia hedleyi* (Fulton, 1907) | MV1082 | Cairns, QLD |
| *Falspleuroxia overlanderensis* Solem, 1997 | WAM S70235 | Shark Bay, WA |
| *Figuladra incei curtisiana* (Pfeiffer, 1864) | NMV F219323 | Mt Archer, QLD |
| *Gnarosophia bellendenkerensis* (Brazier, 1875) | NMV F226513 | Alligator creek, QLD |
| *Hadra bipartita* (Férussac, 1823) | AM C.476663 | Green Island, QLD |
| *Kimboraga micromphala* (Gude, 1907) | AM C.463554 | Windjana Gorge, WA |
| *Kymatobaudinia carrboydensis* Criscione & Köhler, 2013 | WAM 49172 | Carr Boyd Ranges, WA |
| *Marilynessa yulei* (Forbes, 1851) | MV1265 | Brandy Creek, QLD |
| *Mesodontrachia fitzroyana* Solem, 1985 | AM C.476985 | Victoria River District, NT |
| *Nannochloritis layardi* (Gude, 1906) | AM C.477826 | Somerset, QLD |
| *Neveritis poorei* (Gude, 1907) | MV1054 | Mt Elliot, QLD |
| *Noctepuna mayana* (Hedley, 1899) | AM C.478270 | Diwan, QLD |
| *Ordtrachia australis* Solem, 1984 | AM C.462736 | Victoria River District, NT |
| *Patrubella buxtoni* (Brazier, 1880) | AM C.478884 | Moa Is., Torres Strait |
| *Plectorhagada plectilis* (Benson, 1853) | WAM S70240 | Shark Bay, WA |
| *Rhynchotrochus macgillivrayi* (Forbes, 1851) | AM C.478271 | Diwan, QLD |
| *Semotrachia basedowi* (Hedley, 1905) | AM C.476884 | Musgrave Ranges, WA |
| *Sinumelon vagente* Iredale, 1939 | WA 61253 | Mt Gibson, WA |
| *Sphaerospira fraseri* (Griffith & Pidgeon, 1833) | MV1104 | Benarkin State Forest, QLD |
| *Tatemelon musgum* (Iredale, 1937) | AM C.476881 | Musgrave Ranges, WA |
| *Tolgachloritis jacksoni* (Hedley, 1912) | NMV F226521 | Mt Garnet, QLD |
| *Torresitrachia torresiana* (Hombron & Jacquinot, 1841) | AM C.477860 | Weipa, Cape York Peninsula, QLD |

\*All localities within Australia unless otherwise indicated

We used FastUniq v1.1 (Xu *et al*. 2012) to remove duplicates, and Trimmomatic v0.22 (Bolger *et al*. 2014) to trim and remove low quality reads and adaptor sequences (minimum average quality score threshold of 20 per 8nt window). Reads shorter than 40 bases after trimming were discarded. The trimmed reads were then mapped onto the transcriptome sequences used for the probe design using BFAST v0.7.0a (Homer *et al*. 2009) with a single index of 22nt without mismatch. After creating pileup files using Samtools v0.1.19 (Li et al. 2009), VarScan v2.3.7 (Koboldt *et al*. 2012) was used to call variants and produce a final consensus sequence for each taxon per exon. Viewing the initial BAM alignments showed that exon boundaries were often not conserved between *L.gigantea* and the Camaenidae. In these cases (Figure S5) the reference exons were split to reflect the actual exon boundaries in the Camaenidae. The reads where then mapped to the revised exon reference and consensus sequences made as outlined above. To flag potential pseudogenes and paralogs we identified consensus sequences with an elevated proportion of variable sites (> 3% heterozygote sites) and reviewed the corresponding read alignments (BAM files) using the Integrative Genomics Viewer (IGV: Thorvaldsdóttir *et al*. 2013). All sequences with greater than 3% ambiguous sites where removed from the final dataset. Exons where more than 10% of the taxa contained greater than 3% ambiguous sites were discarded entirely.

We again used TreSpEx to assess conflicting phylogenetic signal. We screened for hidden paralogs based on five *a priori* phylogenetic hypotheses representing well supported clades (≥ 75% bootstrap support) within the Australasian camaenid radiation as delineated by Hugall and Stanisic (2011), namely the Hadroid group (clade 1 – 4 inclusive), the far-northern (sister clades 5 and 6) and north-eastern (clade 7) Chloritid groups, a group dominated by arid and monsoonal camaenids (clade 11) previously recognised as the subfamily Sinumeloninae (e.g. Solem 1992), and a phenotypically and ecologically diverse group dominated by eastern Australian wet forest taxa (sister clades 8 and 9). Gene trees for each of the 490 genes (exons from the same gene were combined as one partition) were constructed using the GTRGAMMA model and 100 fast bootstraps in RAxML (Stamatakis 2006). TreSpEx was run using the same settings as the analysis for the transcriptome dataset (i.e. TreSpEx considered nodes for strong conflict, long branches, and short branches in that order with parameters upbl and lowbl set to 5 and blt 0.00001).

### Comparison to the Agalma pipeline

As an independent qualification of the manually curated 500 gene set we ran the fully automated orthology determination pipeline Agalma (Dunn *et al*. 2013) (Figure 1). We commenced this pipeline from the ‘postassemble’ step which first identified open reading frames and putative coding regions (Dunn *et al*. 2013). Homolog clusters were then identified using an all-by-all tblastx, followed by clustering using the Markov Clustering algorithm (MCL) (Figure 1). Homolog clusters were then translated and aligned using MAFFT (Katoh & Standley 2013) and gene trees estimated using RAxML. To identify orthologous sequences, the genealogies were then screened for ‘optimally inclusive subtrees’ which contain only a single representative of each species. Multiple orthologous subtrees can be delineated per homolog cluster, potentially allowing paralogs to be separated and retained. The surviving subtrees were filtered based on the number of taxa (set to greater than four taxa) and realigned for subsequent phylogenetic analysis. We then identified Agalma homologous clusters that corresponded to the manually curated 500 gene set using BLAST (blastp, e-value cut off of 1e-10).

### Phylogenetic analysis

After removal of paralogs or sequences with excessive polymorphism (>3% dissimilarity), our phylogenomic datasets were refined by removing any regions of ambiguous alignment through the use of Gblocks (Castresana 2000) (which is built into the Agalma pipeline) and manual masking. We reconstructed maximum likelihood trees using the program RAxML (Stamatakis 2006) for datasets resulting from both the manual curation and the Agalma pipeline. PartitionFinder (Lanfear *et al*. 2012, 2014) was used to identify suitable models and partitioning schemes, implemented with 1% heuristic r-cluster searches, optimized weighting, RAxML likelihood calculations, and model selection based on BIC scores. In all cases, nodal support was assessed by performing 100 full non-parametric bootstraps.

We analysed two datasets resulting from the Agalma pipeline. The first dataset comprised ortholog clusters that corresponded to the manually curated 500 gene set (here on referred to as the ‘Agalma equivalent dataset’). The second dataset consisted of all ortholog clusters which had high taxon coverage ≥18), and were derived from homolog clusters containing only a single ortholog cluster (from here on referred to as the ‘Agalma best dataset’); that is, Agalma homolog clusters containing multiple copies, albeit diagnosable, were not considered further. Finally, we reconstructed a phylogeny for the camaenid dataset obtained through exon-capture and included sequences from the five camaenid transcriptomes presented herein, as well as sequences of *Cornu aspersum* as an outgroup.

## Results

### Transcriptome assembly and homolog clustering

The number of paired reads obtained for each of the 21 eupulmonate species sequenced ranged from 7.8M to 31.6M (Table 3). Trimming and de novo assembly statistics are presented in Table 3. The number of *L. gigantea* reference genes with BLAST matches ranged from 7,011 to 9,699 per assembly (Table 3), 5,490 of which had homologous sequences in at least 18 of the 21 transcriptome assemblies.

**Table 3.** Summary statistics for sequencing and *de novo* assembly of 21 eupulmonate transcriptomes

| Species | Pairs of raw reads | Proportion of reads after trimming | Trinity contigs | BLAST hits 1e-10 (*L. gigantea*) | *L. gigantea* genes with hits | No. of the 500 single copy genes |
| --- | --- | --- | --- | --- | --- | --- |
| *Ramogenia challengerii* | 11,726,377 | 0.84 | 103,471 | 14,665 | 7,011 | 488 |
| *Austrochloritis kosciuszkoensis* | 11,357,080 | 0.85 | 107,810 | 16,238 | 7,522 | 495 |
| *Sphaerospira fraseri* | 31,594,841 | 0.85 | 179,695 | 23,910 | 9,433 | 500 |
| *Thersites novaehollandiae* | 15,620,892 | 0.85 | 118,298 | 17,330 | 7,869 | 492 |
| *Chloritobadistes victoriae* | 26,433,009 | 0.85 | 148,817 | 20,453 | 8,792 | 498 |
| *Amimopina macleayi* | 7,874,195 | 0.97 | 93,250 | 17,258 | 8,091 | 494 |
| *Cochlicopa lubrica* | 8,074,560 | 0.97 | 111,396 | 21,675 | 9,086 | 497 |
| *Asperitas stuartiae* | 9,322,853 | 0.97 | 104,942 | 15,491 | 7,460 | 491 |
| *Cassidula angulifera* | 14,281,906 | 0.97 | 105,803 | 16,981 | 8,083 | 489 |
| *Apoecus cf apertus* | 9,362,182 | 0.97 | 119,711 | 21,275 | 9,095 | 497 |
| *Fastosarion cf virens* | 14,904,669 | 0.84 | 127,454 | 18,306 | 7,987 | 494 |
| *Cornu aspersum* | 21,273,910 | 0.86 | 160,490 | 23,114 | 9,254 | 498 |
| *Limax flavus* | 14,907,395 | 0.84 | 116,088 | 19,071 | 8,349 | 497 |
| *Lamprocystis* sp. | 22,539,699 | 0.97 | 128,611 | 23,797 | 9,679 | 499 |
| *Milax gagates* | 11,263,950 | 0.97 | 92,337 | 16,541 | 7,041 | 490 |
| *Oxychilus alliarius* | 12,925,111 | 0.97 | 136,044 | 21,183 | 8,940 | 499 |
| *Terrycarlessia turbinata* | 16,985,068 | 0.84 | 141,421 | 17,073 | 7,778 | 489 |
| *Victaphanta atramentaria* | 11,312,274 | 0.86 | 101,127 | 16,584 | 7,466 | 490 |
| *Austrorhytida capillacea* | 10,154,817 | 0.96 | 88,525 | 15,352 | 7,118 | 477 |
| *Smeagol phillipensis* | 6,393,571 | 0.96 | 95,429 | 23,067 | 9,699 | 497 |
| *Semperula maculata* | 12,461,924 | 0.97 | 76,847 | 21,851 | 9,276 | 492 |

Of the 288 genes used in a previous molluscan phylogenomic study (Kocot *et al*. 2011), 130 were single-copy for all eupulmonates considered here, while 146 contained paralogs in at least one species (mean p-distance between paralogs within a sample was 0.28, ranging from 0.16-0.46). We could not unambiguously qualify the remaining 12 genes from the Kocot *et al*. study as they were poorly represented in our transcriptomes. Prioritising genes with high taxon coverage and long exon length, we assessed additional alignments of candidate homolog clusters until we reached a total of 500 single-copy genes. In addition to the 146 Kocot genes shown to be paralogous within the eupulmonates, we identified and qualified 62 multi-copy genes during the course of this work. The resulting manually curated 500 single-copy gene set is 98.5% taxa complete (i.e. sequence present for each gene and taxon) and 93.1% character complete (Figure 4d), with an average gene length of 1,190nt, ranging from 228nt to 6,261nt. In total, the final alignment of this gene set represents 512,958nt. Approximately 12% of the sequences in the final gene-by-species matrix were derived by merging fragmented transcripts.

Based on the all-by-all BLAST comparison of the *L. gigantea* genes, 347 of our final 500 genes had a single hit at an e-value threshold of 1e-10 (i.e. single copy status was consistent between the *L. gigantea* reference and the eupulmonates), while the remainder had multiple hits, indicative of the presence of close paralogs in the reference. Conversely, of the 208 genes qualified as multiple-copy for the eupulmonates (146 from the Kocot gene set plus 62 from this study), 134 only had one hit within the *L. gigantea* gene set (i.e. just over half of the multiple-copy gene set are potentially single copy for patellogastropods). These results broadly correspond to the orthology designation in the OMA (Orthologous MAtrix) database.

Across the 500 single-copy genes, the p-distance between the two rhytidids, *Terrycarlessia turbinata* and *Victaphanta atramentaria*, ranged from 0.02 to 0.13 (average of 0.06; Figure 3). This family is thought to have originated 120 Mya (Bruggen 1980; Upchurch 2008). However, the Australian rhytidids probably represent a more recent radiation (Herbert et al. 2015, Moussalli and Herbert 2016). Similarly, p-distance between the two camaenids, *Sphaerospira fraseri* and *Austrochloritis kosciuszkoensis*, ranged from 0.01 to 0.13 (average of 0.04). This group is thought to have originated in the Oligo-Miocene approximately 30 Mya (Hugall & Stanisic 2011). All genes had a higher relative substitution rate than the commonly used phylogenetic marker 28S, and were on average approximately four times slower than COI (Figure 3).

### Qualification of orthology using gene tree assessments

TreSpEx analyses of all 500 genes found no well supported conflict with the *a priori* phylogenetic hypotheses, suggesting that hidden paralogs (i.e., genes represented by a single sequence per taxon yet paralogous across multiple taxa) were absent from our dataset. Furthermore, this analysis also showed no evidence of cross sample contamination, nor any evidence of suspect long internal branches within the Stylommatophora.

### Qualification of orthology using exon-capture

We enriched and sequenced all 1,646 targeted exons, from 490 genes, when considering all 22 samples collectively. We first mapped reads to the original reference used in the probe design with exon boundaries delineated based on the *L. gigantea* genome. Examination of the resulting read alignments (BAM files) identified 437 exons which contained multiple internal exon boundaries within the Camaenidae (Figure S4). Accordingly, the mapping reference was modified to account for exon-splitting (including the removal of 163 exons that were shorter than 40 bp after splitting), with the final revised reference comprising 2,648 exons representing 417,846nt (Supplementary Table 1). We targeted an average of five exons per gene.

We then remapped reads to the revised reference (coverage and specificity statistics presented in Table 4) and flagged resulting consensus sequences which exhibited elevated polymorphism (> 3% heterozygote sites). There were 508 exons where at least one taxon exhibited elevated polymorphism. Of these, 105 exons had greater than 10% of the taxa (typically two or more taxa, taking into account missing taxa) exhibiting elevated polymorphism. Based on an examination of the corresponding read alignments, 95 exons were classified as having lineage specific pseudogenes or paralogs, four contained evidence of processed pseudogenes, and six where the alignment was complicated by the mapping of unrelated reads containing small, highly similar domains (see Figure S4-S8 for examples of each case). These 105 exons were removed prior to phylogenetic analyses. For the remaining 403 exons only the consensus sequences for the taxa with elevated polymorphism were removed from the final alignment. In total, 3.7% of the sequences were removed from final data matrix due to elevated polymorphism. The final exon capture data matrix was 98% taxa complete and 95% character complete.

**Table 4.** Sequencing and mapping summary statistics for the exon capture experiment.

| Species | No. raw paired end reads | Proportion of pairs of reads retained after duplicate removal | Proportion retained after Trimmomatic | Proportion of reads mapped to the final reference | Average coverage per exon | Proportion of exons captured (total 2648 exons) |
| --- | --- | --- | --- | --- | --- | --- |
| *Boriogenia hedleyi* | 836,437 | 0.60 | 0.97 | 0.64 | 145 | 0.96 |
| *Falspleuroxia overlanderensis* | 170,769 | 0.69 | 0.98 | 0.74 | 41 | 0.88 |
| *Figuladra incei curtisiana* | 1,117,954 | 0.57 | 0.96 | 0.6 | 167 | 0.97 |
| *Gnarosophia bellendenkerensis* | 1,490,686 | 0.57 | 0.98 | 0.63 | 235 | 0.98 |
| *Hadra bipartita* | 659,509 | 0.6 | 0.98 | 0.7 | 131 | 0.96 |
| *Kimboraga micromphala* | 186,942 | 0.86 | 0.99 | 0.73 | 55 | 0.90 |
| *Kymatobaudinia carrboydensis* | 666,965 | 0.78 | 0.98 | 0.63 | 145 | 0.94 |
| *Marilynessa yulei* | 865,712 | 0.56 | 0.97 | 0.62 | 139 | 0.97 |
| *Mesodontrachia fitzroyana* | 429,572 | 0.85 | 0.98 | 0.61 | 102 | 0.91 |
| *Nannochloritis layardi* | 179,432 | 0.86 | 0.97 | 0.72 | 50 | 0.90 |
| *Neveritis poorei* | 1,313,049 | 0.57 | 0.96 | 0.62 | 205 | 0.95 |
| *Noctepuna mayana* | 297,503 | 0.77 | 0.98 | 0.73 | 81 | 0.93 |
| *Ordtrachia australis* | 670,743 | 0.65 | 0.94 | 0.86 | 222 | 0.92 |
| *Patrubella buxtoni* | 492,474 | 0.82 | 0.97 | 0.7 | 125 | 0.92 |
| *Plectorhagada plectilis* | 220,636 | 0.81 | 0.98 | 0.76 | 65 | 0.90 |
| *Rhynchotrochus macgillivrayi* | 340,338 | 0.85 | 0.98 | 0.7 | 96 | 0.92 |
| *Semotrachia basedowi* | 290,966 | 0.92 | 0.88 | 0.83 | 119 | 0.92 |
| *Sinumelon vagente* | 282,838 | 0.86 | 0.97 | 0.75 | 86 | 0.92 |
| *Sphaerospira fraseri* | 796,591 | 0.56 | 0.98 | 0.66 | 130 | 0.98 |
| *Tatemelon musgum* | 242,614 | 0.87 | 0.99 | 0.7 | 66 | 0.91 |
| *Tolgachloritis jacksoni* | 1,207,039 | 0.38 | 0.97 | 0.65 | 139 | 0.95 |
| *Torresitrachia torresiana* | 192,031 | 0.87 | 0.98 | 0.74 | 61 | 0.90 |

Based on the TreSpEx analyses, four genes did not support the monophyly of the ‘Far North Chloritid’ group, but rather placed (*Nannochloritis layardi* and *Patrubella buxtoni*) as sister to the ‘North-East Chloritid’ group (Figure 5). We concluded that this was not the result of hidden paralogy, but rather due to insufficient lineage sorting of relatively conserved genes. An additional five genes were in conflict with the *a priori* taxonomic hypotheses, however, these represented cases where the genes were small and the proportion of phylogenetically informative sites was low. Five genes were flagged as having at least one internal branch which was greater than five times the average. Assessment of the alignments and corresponding genealogies indicated that they represented deep basal divergence between well supported major clades, and was not reflective of hidden paralogy.

**Figure 5.**
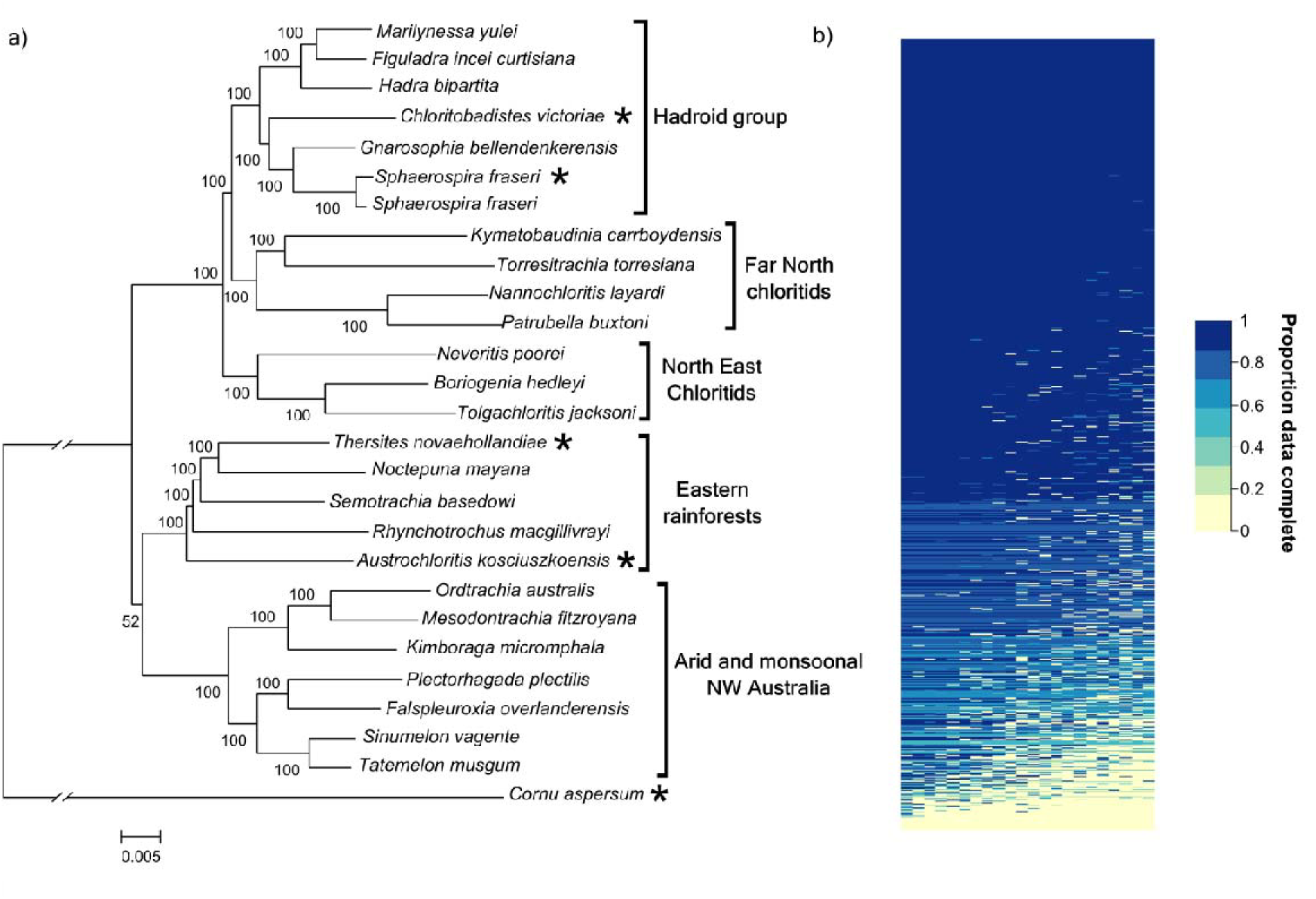
Maximum likelihood phylogeny of 26 Australian camaenid land snails. (a) Phylogenetic reconstruction based on nucleotides sequences from 2,648 exons obtained through exon-capture. Sequences for the taxa marked with asterisks were derived from transcriptome datasets. Numbers on branches indicate bootstrap nodal support. (b) Heat map showing the proportion of available sequences for each sample per gene (sorted left to right by proportion of data present per sample; top to bottom by proportion of data present per exon).

Finally, we enriched another representative of *Sphaerospira fraseri*, one of the reference species used in the probe design. Comparing the mapped consensus genomic sequence to the transcriptome reference we found only minor mismatch, reflective of intraspecific variation as the two samples came from different populations (the exons had a median p-distance of 0.8%). Furthermore, for this species at least, all reference genes constructed from multiple transcript fragments were consistent with those captured from genomic DNA (i.e. chimeras of unrelated fragments were not created) and showed no evidence of paralogy or elevated heterozygosity.

### Comparison to Agalma pipeline

Using the Agalma pipeline we identified 11,140 ortholog clusters. Of these ortholog clusters 635 corresponded to 457 of our 500 single-copy gene set. We refer to this dataset as the “Agalma equivalent” dataset, and is 61% taxa complete and 54% character complete. Many of the genes were represented by multiple ortholog clusters in the Agalma analysis, many of which contained fewer taxa relative to that obtained via manual curation (Figure 2). Rather than paralogs, in all cases fragmentation in the transcriptome assemblies resulted in the splitting of homolog clusters into multiple ortholog clusters, each representing the same locus but containing a different subset of taxa (see example in Figure S3). Of the 43 single-copy genes not picked up by Agalma, five were not annotated in the ‘postassemble’ step, 12 were annotated but not recovered by the all-by-all BLAST, 18 were recovered by the all-by-all BLAST but dropped during the clustering step, and eight made it to the initial clusters but failed the alignment and trimming step prior to the gene tree reconstruction. Failure to recover these genes during the BLAST comparison, clustering and alignment steps is most likely due to a combination of frameshift errors and transcript fragmentation, and in certain cases, resulting in the taxon sampling threshold and cluster size criteria not being met.

Of the 11,140 ortholog clusters there were 546 clusters that contained sequences of at least 18 taxa and that had one ortholog cluster per homolog cluster. Of these, 171 were also contained in our 500 single-copy gene set. Hence, the Agalma pipeline identified 375 genes in addition to the 500 manually curated genes, which had optimum taxon sampling. The majority of these genes also represented the full CDS with 89% representing at least 80% of the length of the respective *L. gigantea* gene. We refer to this dataset as the “Agalma best” dataset and is 92% taxa complete and 85% character complete.

### Phylogenetic analysis

We reconstructed phylogenies from three ortholog datasets for comparison: (1) the manually curated 500 single-copy gene set (Figure 4a, d),(2) the Agalma equivalent dataset consisting of 635 orthologous clusters which corresponded to 457 of the 500 single-copy genes (Figure 4b, e), and (3) the Agalma best dataset consisting of 546 orthologous clusters which had 18 or more taxa and were the only orthologous cluster from the respective homolog cluster (Figure 4c, f). Of the manual curated dataset, 1.6% of the alignment was removed by Gblocks prior to phylogenetic analysis. The phylogenies for the 500 single-copy gene set and the Agalma best dataset had identical topologies, supporting all major clades with very high bootstrap support, namely Helicoidea, Limacoidea, Orthurethra, the Australian rhytidids and the Stylommatophora (Figure 4a, c). In terms of phylogenetic relationships, the Rhytididae forms a sister relationship with the Limacoidea, and the Helicoidea occupies a basal position within Stylommatophora. In contrast, while also supporting the monophyly of all major clades, the phylogeny based on the ‘Agalma equivalent’ dataset places Orthurethra in a basal position within Stylommatophora, (Figure 4b).

Of the Camaenidae exon capture dataset, 5% of the alignment was removed by Gblocks prior to phylogenetic analysis. The resulting phylogeny supported all major groups previously recognised by Hugall and Stanisic (2011). In terms of phylogenetic relationships, the two Chloritid groups formed a clade with the Hadroid group, with the Far-northern chloritids sister to the hadroids. There was poor resolution regarding the phylogenetic positions of the two remaining groups, the Eastern rainforests and the arid and monsoonal NW Australian clades (Figure 5).

## Discussion

The identification and qualification of orthology is a critical prerequisite for sound phylogenetic inference. Our approach of orthology assessment involved an initial assessment and manual editing of homolog clusters, allowing us to correct for multiple isoforms and errors such as sequence fragmentation, frame-shifts and mis-indexing. Using this approach, we qualified the orthology and single-copy status of 500 genes across the eupulmonates, 130 of which were used in a previous phylogenomic study of the Mollusca (Kocot *et al*. 2011). The resulting 500 gene data matrix is the most complete produced for a major molluscan lineage to date, both in terms of taxon and character completeness. We further qualified orthology by capturing and sequencing 490 of the 500 genes from genomic DNA, revealing the presence of paralogs and/or pseudogenes otherwise not evident from the transcriptome data. Although the automated pipeline Agalma recovered the majority of the 500 genes as single copy and identified 375 additional putatively orthologous genes for the eupulmonates, it was hampered by transcript fragmentation within the assemblies. Furthermore, supported topologies for the 21 eupulmonate species were not entirely consistent between the manually curated and Agalma equivalent dataset, potentially a consequence of lower data matrix completeness in the latter. We discuss approaches to ortholog determination and implications for phylogenetic inference below.

### Ortholog determination

To date, most transcriptome based phylogenomic studies have focused on resolving relatively deep evolutionary relationships (e.g. Kocot *et al*. 2011; Smith *et al*. 2011; Zapata *et al*. 2014; O’Hara *et al*. 2014; Misof *et al*. 2014), and a number have relied on annotated ortholog databases for the initial screening of suitable genes, such as OMA (Altenhoff *et al*. 2015), OrthoDB (Waterhouse *et al*. 2013), and the ortholog dataset associated with HaMStR (Ebersberger *et al*. 2009). Such databases are typically limited in the number of representatives per lineage (e.g., Tatusov *et al*. 2003; Ranwez *et al*. 2007; Waterhouse *et al*. 2013; Altenhoff *et al*. 2015). Nevertheless, it is a reasonable assumption that orthologous genes qualified as single-copy across many highly divergent taxa are more likely to maintain single-copy status with greater taxonomic sampling. We tested this idea at a preliminary stage of our work by first assessing genes used in a phylogenomic study of the Mollusca (Kocot *et al*. 2011). In that study, orthologous genes were identified using the program HaMStR, based on a 1,032 ortholog set resulting from the Inparanoid orthology database (Ostlund *et al*. 2010). We found that just under half of the genes used in Kocot *et al*. (2011) were paralogous within the eupulmonates. To some extent the high proportion of the Kocot *et al*. gene set being paralogous is due to the limited representation of eupulmonates in that study, and for these few taxa paralogs may have been absent. Alternatively, in such deep phylogenomic studies lineage-specific duplication may have manifested as in-paralogs and were dealt with by retaining one copy from the in-paralog set at random (Kocot *et al*. 2011; Dunn *et al*. 2013) or based on sequence similarity (Ebersberger *et al*. 2009). However, with an increase in taxonomic sampling, such paralogy may extend across multiple taxa and, unless conservation of function can be established (i.e. isorthology, Fitch 2000), these genes would no longer be suitable for phylogenetic analysis.

When the 500 gene set was compared to the OMA database (Altenhoff *et al*. 2015), which at the time of this analysis only incorporated a single molluscan genome, namely *L. gigantea*, we found a similarly high proportion of eupulmonate specific paralogy. A more interesting result arising from this comparison, however, was that many genes classified as having putative paralogs in *L. gigantea* were single-copy across the eupulmonates. We cannot ascertain at this stage whether this is a consequence of duplication being derived within Patellogastropoda, the lineage containing *L. gigantea,* or the consequence of duplicate loss in the ancestral eupulmonate. Nevertheless, this result highlights that potentially suitable genes may be overlooked when restricted to ortholog database designations, especially when such databases have poor representation of the relevant lineage. Accordingly, although we used the *L. gigantea* gene set as a reference with which to identify and cluster homologous sequences, we did not rely on orthology database designations of the *L. gigantea* gene set to guide which genes to consider when assessing orthology across the eupulmonates examined here.

### Automated vs manually curated aided pipelines

Pipelines that fully automate homology searches and clustering, orthology qualification, and final alignments are highly desirable for efficiency, consistency, and repeatability. Moreover, reference free methods, like that implemented in Agalma, are also highly desirable in cases where the study taxa are poorly represented in ortholog databases. There are characteristics of assembled transcriptome sequences, however, that can challenge fully automated methods, including transcript fragmentation, mis-indexing, frameshifts and contamination, and these aspects necessitate careful manual appraisal and editing (Philippe *et al*. 2011; O’Hara *et al*. 2014). Although recent phylogenomic studies have, to varying degrees, incorporated manual appraisal, such checks are typically conducted at the final proofing stage (e.g. Kocot *et al*. 2011; Simmons & Goloboff 2014). In this study, we purposefully addressed the abovementioned issues at an early stage following the initial alignment of homologous sequences. The most important aspect of our manual curation was the creation of consensus sequences from fragmented transcripts (see also: O’Hara *et al*.2014), which in turn ensured maximum retention of data (particularly for probe design) and placed subsequent orthology assessment on a sounder footing. Consequently, our final data matrix was highly complete (93% character complete whereas the ‘Agalma best’ dataset was 85% character complete).

The Agalma analysis confirmed the single-copy, orthology status for the majority of the 500 manually curated gene set, but it was hampered by transcript fragmentation within the transcriptome assemblies. In all cases where multiple ortholog clusters were derived using Agalma for any one of our 500 genes, this was due to transcript fragmentation, not missed paralogy. In essence, alignments of fragmented transcripts (whether or not they were partially overlapping) resulted in poorly reconstructed gene trees, which in turn misled subsequent tree pruning and ortholog clustering (e.g. Figure S3). Consequently, for the ‘Agalma equivalent’ dataset, both taxon and character completeness was poor relative to the manually curated data matrix. To our knowledge, no fully automated phylogenomics pipeline currently implements the consensus of fragmented sequences, and studies that have made the effort to retain multiple fragments, as in this study, have decided which sequences to retain and merge manually (e.g., Rothfels *et al*. 2013; O’Hara *et al*. 2014). The issue of working with fragmented assemblies can be addressed, however, by incorporating an automated consensus making algorithm such as TGICL (Pertea *et al*. 2003) into the pipeline to address fragmentation at the homolog alignment stage. Doing so is particularly desirable, given that manual curation of homologous sequences requires considerable time investment.

A major strength of automated pipelines is that they enable a more comprehensive screening of putative orthologous genes. Manual curation requires considerable effort, and while more candidate genes were identified than were assessed, we ceased the manual assessment once our target of 500 genes had been attained. The Agalma analyses had no constraints, however, hence all possible orthologous clusters were considered. Consequently, we identified an additional 375 ortholog clusters which met a strict taxa completeness threshold (18 taxa or more) and represented the only ortholog cluster arising from original homolog clusters. These genes (i.e. the ‘Agalma best’ dataset) reconstructed a phylogeny that was very similar to the manually curated dataset. While beyond the scope of this study, there is potential for these genes to be included in future probe designs and further qualification of these additional genes using exon-capture (see below) would be highly desirable.

### Phylogenetic inference

The 500 gene set represents a significant contribution towards advancing molecular phylogenetics of the eupulmonates, providing the capacity to resolve both evolutionary relationships at shallow to moderate depths, and deep basal relationships. The phylogenetic reconstructions presented here are well resolved and support the *a priori* taxonomic hypotheses used as part of the orthology assessment. In terms of deeper relationships, reconstructions based on the two most complete datasets are consistent, namely the monophyly of Stylommatophora, within which Helicoidea is basal, and the sister relationship between the Rhytidoidea and the Limacoidea. For the less complete Agalma equivalent dataset, however, Orthurethra is basal within Stylommatophora, albeit with marginal support. Without greater taxonomic sampling of all the major lineages within the eupulmonates, however, a comprehensive phylogenetic assessment is beyond the scope of this study. Nevertheless, these phylogenomic datasets do afford greater resolution of deeper relationships than obtained in previous molecular studies (Wade *et al*. 2001, 2006). Secondly, convergence in supported topology between the two most complete and largely independent datasets (only 171 genes were in common), and the inconsistency between the manually curated and Agalma equivalent dataset (sharing 458 genes), suggests the possible importance of data matrix completeness in resolving short, basal internodes.

### Exon-capture

One of the overarching objectives of this study was to identify and qualify 500 genes suitable for exon-capture work within the eupulmonates. Here we sequenced and analysed a small dataset for the family Camaenidae principally as a means to further qualify orthology. There are two principle outcomes from this exploration. First, for all reference sequences based on the concatenation of fragmented transcripts, there was no evidence that erroneous chimeric sequences were created. Second, as was the case with the increased sampling in the transcriptome work, the pervasiveness of lineage-specific duplication was also evident from the exon-capture experiment. Despite qualification of single-copy orthology of the transcriptome dataset, increased taxonomic sampling within the family Camaenidae revealed lineage-specific duplication for potentially as high as one fifth of the targeted exons. In the great majority of cases, however, a very small proportion of taxa exhibited putative paralogy or pseudogenes, and removal of the affected exon per taxon only reduced the completeness of the final dataset by 3.7%. Similar results were achieved for the brittle stars with 1.5% of their target discarded due to putative paralogs or pseudogenes (Hugall *et al*. 2015). It is possible that these putative paralogs were only detected in the genomic sequencing because they were not expressed in the transcriptomes.

Within the Australian Camaenidae, uncorrected distances for the majority of the genes did not exceed 13%. This level of sequence variability is within the range of mismatch that is tolerated by in-solution exon-capture protocols (Bi *et al*. 2012; Bragg *et al*. 2015; Hugall *et al*. 2015). This was qualified here given the high proportion of target recovery (>95%) across a broad representation of the camaenid diversity. As was the case for the Euplumonata phylogeny presented above, our preliminary phylogenomic dataset for camaenids provides considerable resolution, particularly among the chloritis and hadroid groups which to date have been difficult to resolve (Hugall & Stanisic 2011).

Expanding the bait design to enrich across the Australasian camaenid radiation, indeed the family Helicoidea, would require the incorporation of multiple divergent reference taxa into the bait design. Recent “anchored enrichment” approaches to bait design (e.g. Lemmon *et al*. 2012; Faircloth *et al*. 2012) target highly conserved regions to allow capture across highly divergent taxa. By contrast, the approach taken here is to target both conserved and highly variable regions, and where possible the full coding region (Bi *et al*. 2012; Bragg *et al*. 2015; Hugall *et al*. 2015). Accordingly, this would require substantially greater reference diversity to be incorporated into the bait design relative to the anchored approach to capture across highly divergent lineages (e.g. across families). Recently, Hugall *et al*. (2015) used a similar approach to the one in the present study, but designed baits based on ancestral sequences, rather than representative tip taxa, to reduce the overall size of the reference set. Using this approach, Hugall *et al*. successfully enriched and sequenced both conserved and highly variable exons across the entire echinoderm class Ophiuroidea, spanning approximately 260 million years. Here we have presented a simple bait design targeting a specific family, but our transcriptome dataset could be used to produce a more diverse bait design to facilitate a more comprehensive study of Eupulmonata phylogenetics and systematics.

## Acknowledgements

We thank Andrew Hugall for bioinformatics advice and Felipe Zapata for assistance with running Agalma; Devi Stuart-Fox, Claire Mclean and Mark Phuong for critical feedback on the manuscript; and Dai Herbert for providing images for Figure 4. We also thank the staff at Australian Genome Research Facility (AGRF) and the Georgia Genomics Facility (GGF), specifically Travis Glenn and Roger Neilsen, for guidance and assistance in transcriptome and exon-capture sequencing. LT was supported by Victorian Life Science Computing Initiative (VLSCI) scholarship. The work was supported by a Holsworth Wildlife Research Endowment to LT, an Australian Biological Resources Study (ABRS) grant to FK and AM (RF213-12). This research was undertaken with the assistance of resources from the National Computational Infrastructure (NCI), which is supported by the Australian Government.

## Data Accessibility

Raw high-throughput sequence reads: NCBI Bioproject PRJNA304185

Transcriptome assemblies, gene and exon alignments for the transcriptome analyses, the Camaenidae exon-capture probe set and the data sets used for phylogenetic inference: Dryad (doi:10.5061/dryad.fn627)

## Author Contributions

LCT and AM designed the study. LCT lead the analysis with contribution from AM, TOH, and KDM. LCT, AM and FK collected samples. LCT and AM wrote the manuscript. All authors reviewed and edited the manuscript prior to submission.

